# Parallel analysis of voltage-gated sodium channel subunits reveals preferential colocalizations of beta-1/Nav1.1 and beta-2/Nav1.2

**DOI:** 10.64898/2026.03.12.711489

**Authors:** Tetsushi Yamagata, Toshimitsu Suzuki, Kazuhiro Yamakawa

**Affiliations:** Department of Neurodevelopmental Disorder Genetics, Institute of Brain Science, Nagoya City University Graduate School of Medical Sciences, Nagoya, Aichi 467-8601, Japan; Laboratory for Neurogenetics, RIKEN Center for Brain Science, Wako, Saitama 351-0198, Japan

## Abstract

Voltage-gated sodium channels (VGSCs) are conventionally described as heterotrimers composed of one alpha and two beta subunits. However, the patterns of co-expression of alpha– and beta-subunits in neurons remain unclear. In the present study, we report that alpha– (Nav1.1, Nav1.2, and Nav1.6) and beta– (beta-1 and beta-2) subunits are densely expressed in axon initial segments (AISs) of neurons in the neocortex, hippocampus and cerebellum at postnatal days 14–15 (P14–15) and 8–9 weeks (8–9W). These distributions are largely unique and partially overlapping among brain regions. Notably, in the neocortex and hippocampus, AISs of presumptive parvalbumin-positive inhibitory neurons are positive for Nav1.1 and beta-1, whereas those of excitatory ones are positive for Nav1.2 and beta-2. Similarly, AISs of cerebellar basket cells, which are inhibitory neurons, are positive for Nav1.1 and beta-1, whereas those of granule cells, which are excitatory neurons, are positive for Nav1.2 and beta-2. Nav1.6 is expressed in many of these neurons. Some subunits exhibited distinct distribution patterns at the two postnatal stages analyzed, possibly because of their developmental changes of subcellular localizations. Taken together, these results indicate that combinations of VGSC subunits are largely unique among different neuronal subpopulations. These findings provide a useful reference for understanding the distribution and interactions of VGSC subunits in the brain.

## 1. Introduction

The VGSC family consists of nine alpha (Nav1.1–Nav1.9) and four beta (beta-1–beta-4) subunits in mammals. Four of the nine alphas, Nav1.1, Nav1.2, Nav1.3, and Nav1.6, encoded by *SCN1A*, *SCN2A*, *SCN3A,* and *SCN8A* genes, respectively, are expressed mainly in the central nervous system (CNS) (reviewed in Catterall, 2012). Nav1.1, Nav1.2, and Nav1.6 are expressed predominantly during postnatal stages, whereas Nav1.3 is expressed mostly at embryonic/perinatal stages (Shah et al., 2001; Heighway et al., 2022). We and others reported detailed histological, cellular and subcellular distributions of these alpha subunits and their developmental changes (Ogiwara et al., 2007; Lorincz and Nusser, 2008; Hu et al., 2009; Lorincz and Nusser, 2010; Liao et al., 2010; Ogiwara et al., 2013; Li et al., 2014; Miyazaki et al., 2014; Yamagata et al., 2017; Ogiwara et al., 2018; Ye et al., 2018; Spratt et al., 2019; Yamano et al., 2022; Yamagata et al., 2023; Nelson et al., 2024), in which Nav1.1 and Nav1.2 are rather mutually-exclusive but Nav1.6 is often co-expressed with Nav1.1 or Nav1.2 in multiple brain regions. Nav1.1 is dominantly expressed in neocortical, hippocampal, reticular thalamic, cerebellar Purkinje and basket parvalbumin–positive (PV+) inhibitory neurons but also in excitatory neurons such as thalamic projection neurons, while Nav1.2 is densely expressed in neocortical, hippocampal, cerebellar granule excitatory neurons but also in some inhibitory neurons including striatal projection neurons. The betas, beta-1–beta-4 subunits encoded by *SCN1B*–*SCN4B* genes, respectively, are reported to be expressed in both CNS and peripheral nervous system (O’Malley and Isom, 2015). Although beta-1 (Chen et al., 2004; Wimmer et al., 2015; Kruger et al., 2016; Chen et al., 2023) and beta-2 (Chen et al., 2002; Yu et al., 2003; Miyazaki et al., 2014) are widely expressed at multiple regions of the adult brain, beta-3 is mainly expressed at embryonic/perinatal stages similar to Nav1.3 (Shah et al., 2001) and the expression of beta-4 is highly predominant at the striatum (Miyazaki et al., 2014).

In this study, we investigated histological distributions of Nav1.1, Nav1.2, Nav1.6, beta-1 and beta-2 in the mouse brain at two developmental stages (P14–15 and 8–9W) in a common platform for fair comparisons. From the analysis, we excluded Nav1.3 and beta-3 because of their predominant expressions at embryonic stage and beta-4 because of its rather striatum-restricted expression. We here show that, beta-1 tends to be co-expressed with Nav1.1 in AISs of neocortical, hippocampal, cerebellar Purkinje and basket PV+ inhibitory neurons especially at younger developmental stage, while beta-2 is often co-expressed with Nav1.2 in those excitatory neurons. These findings may refine the VGSC distributions and contribute to the understanding of their interactions and functional cooperation in the brain.

## 2. Materials and Methods

### 2.1. Animal Work Statement

All animal experimental protocols were approved by the Animal Experiment Committees of Nagoya City University (NCU) and the RIKEN Center for Brain Science (CBS). Mice were handled in accordance with the guidelines of these committees.

### 2.2. Antibodies

Rabbit polyclonal antibodies against Nav1.6 (II2; Ogiwara et al., 2007), beta-1, and beta-2 (Wong et al., 2005; Oyama et al., 2006), and goat polyclonal antibodies against Nav1.1 (C-18; sc-16031, Santa Cruz Biotechnology) and Nav1.2 (G-20; sc-31371, Santa Cruz Biotechnology) were used as primary antibodies. The specificities of these antibodies were verified by immunoblotting and/or immunohistochemical analyses as described previously (Wong et al., 2005; Oyama et al., 2006; Ogiwara et al., 2007; Yamagata et al., 2017; Ogiwara et al., 2018).

### 2.3. Immunohistochemistry

Both male and female C57BL/6J mice were used in this study. Immunohistochemical analyses were performed using brains from two to three mice per condition. Sample sizes were determined based on previous studies. Paraffin sections (6-µm thick) were prepared from C57BL/6J mouse brains as described previously (Yamagata et al., 2017; Yamagata et al, 2023). Briefly, mouse brains were fixed with 10mM periodate-75mM lysine-4% paraformaldehyde (PLP) solution (McLean and Nakane, 1974), and the sections were incubated with anti-Nav1.1 (1:500), anti-Nav1.2 (1:500), anti-Nav1.6 (1:500), anti-beta-1 (1:1000), or anti-beta-2 (1:1000) antibodies for 12–15 hours at 4 °C. Biotinylated anti-goat IgG (1:200; BA-9500, Vector Laboratories) or anti-rabbit IgG (1:200; 711-065-152, Jackson ImmunoResearch) antibodies were used as secondary antibodies. Immunoreactivity was visualized using a Vectastain Elite ABC Kit (PK-6100, Vector Laboratories) and developed using a NovaRed Substrate Kit (SK-4800, Vector Laboratories). Images were acquired using a Biozero BZ-8100 microscope (Keyence). Representative images were obtained from two brain sections (N = 1–3).

## 3. Results

### 3.1. Low-magnified investigations of VGSCs in mouse whole brain revealed similar caudal-predominant distributions of Nav1.1 and beta-1, whereas Nav1.2 and beta-2 showed rostral-predominant distributions

We performed a parallel immunohistochemical analysis of Nav1.1, Nav1.2, Nav1.6, beta-1 and beta-2 in the mouse brain. We found that the distribution of beta-1 immunosignals was similar to that of Nav1.1 throughout the brain and predominant in caudal brain regions (Fig. 1A, B). Their expression levels also similarly increased from juvenile stage (postnatal days 14–15: P14–15) to young adult stage (8–9 weeks: 8–9W). On the contrary, distributions of beta-2 and Nav1.2 immunosignals were similar to each other and both rostral-predominant and their expression levels remained the same between the two developmental stages (Fig. 1C, D). Meanwhile, Nav1.6 was widely distributed throughout the brain without apparent regional differences or deviations (Fig. 1E). At 8–9W, the Nav1.6 immunosignals increased compared with those at P14–15. For convenient comparison, these immunohistochemistry images are also presented at a website (https://ndg-ibs.github.io/VGSC_IHC_DB/) in which each figure can be magnified. The first page of the website is shown in Supplementary Fig. S1.

**Figure 1.**
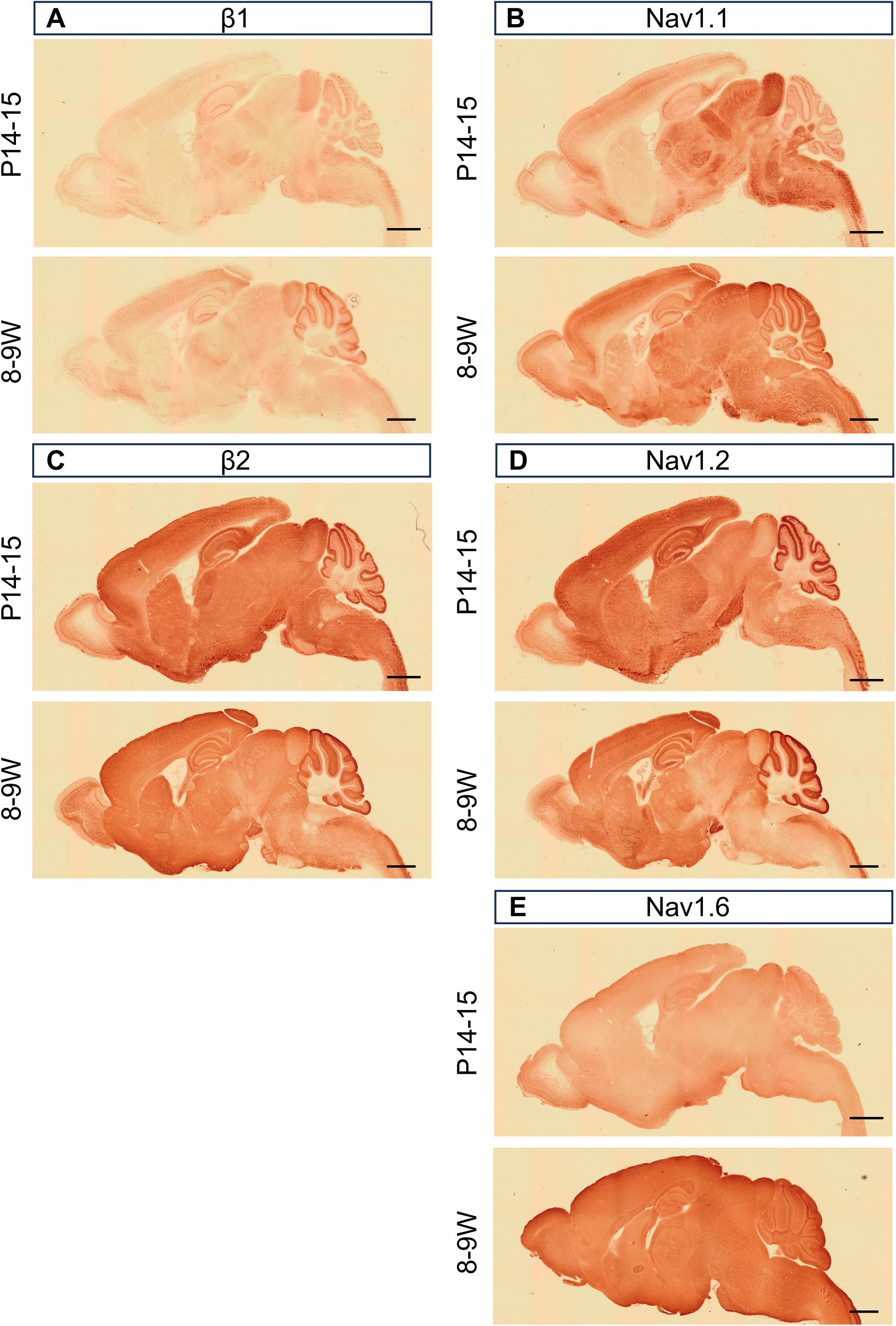
Distinct distributions of Nav1.1, Nav1.2, Nav1.6, beta-1, and beta-2 subunits in the mouse brain. Chromogenic immunostaining for beta-1 (A), Nav1.1 (B), beta-2 (C), Nav1.2 (D), or Nav1.6 (E) was performed on parasagittal sections from C57BL/6J wild-type mouse brains at postnatal days 14–15 (P14–15) and 8–9 weeks (8–9W). Scale bar, 1 mm.

### 3.2. High magnified investigations revealed preferential co-expressions of beta-1 and Nav1.1 in neocortical, hippocampal, and cerebellar inhibitory neurons

We then investigated the immunosignals at higher magnification in neocortex, hippocampus and cerebellum. Because of similar distributions of beta-1 and Nav1.1 at low magnification (Fig. 1A, B), at first we compared these two in parallel at P14–15 and 8–9W (Fig. 2A–D). Although beta-1 immunosignals also appeared in the nuclei in addition to AISs (Fig. 2A1, A2, A5, A6, B1, B2, B5, B6, C1, C2, C5, C6, D1, D2, D5, D6), a previous study using the same anti-beta-1 antibody (Wong et al., 2005) showed that the signals in nuclei are non-specific because they remained in *Scn1b* homozygous knockout mice while the AIS signals were absent in the mice confirming the specificity of the antibody (Wimmer et al., 2015). Similar to the previous study (Wimmer et al., 2015), non-specific nuclear signals in our study were also observed only in excitatory neurons and not in inhibitory neurons.

**Figure 2.**
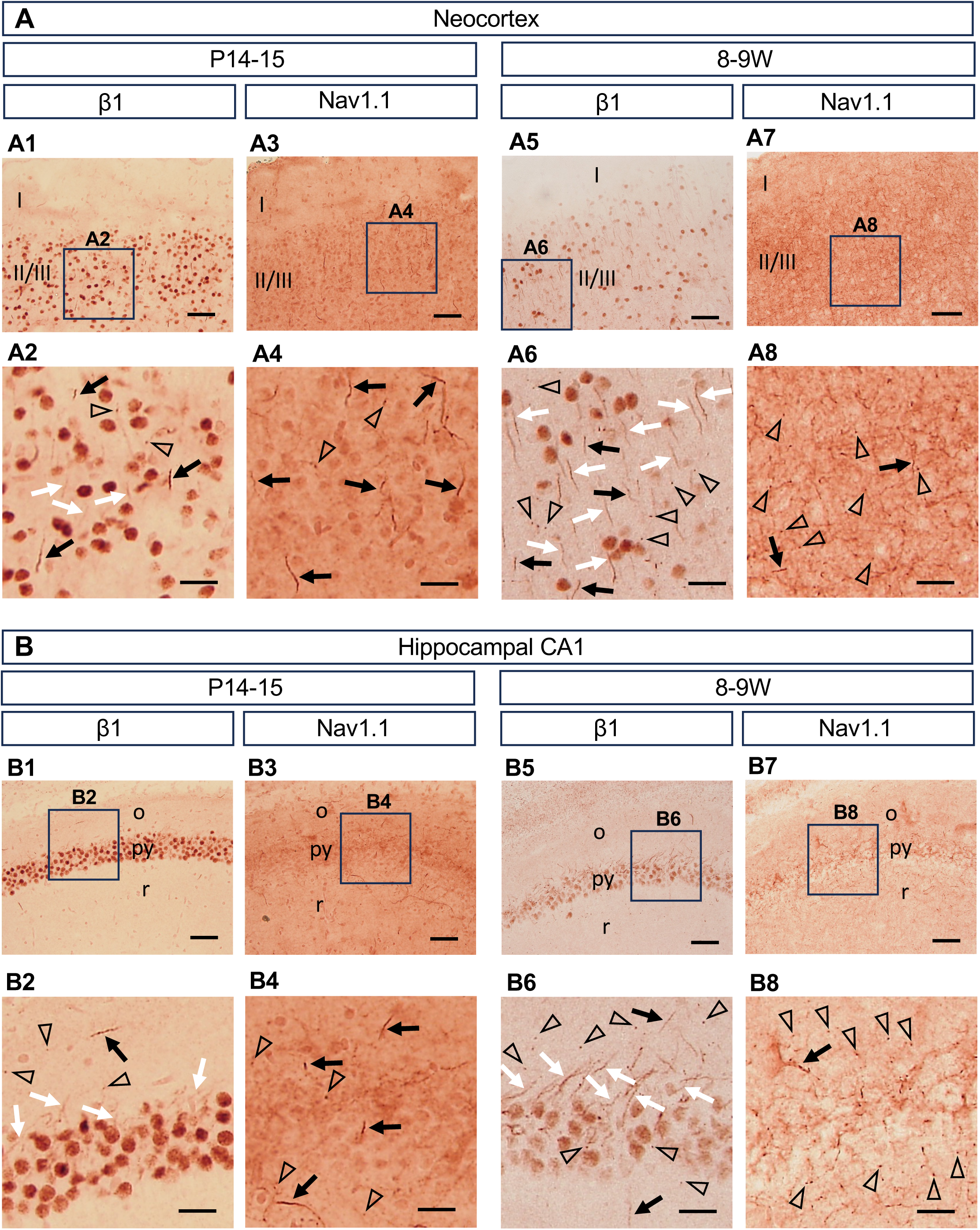
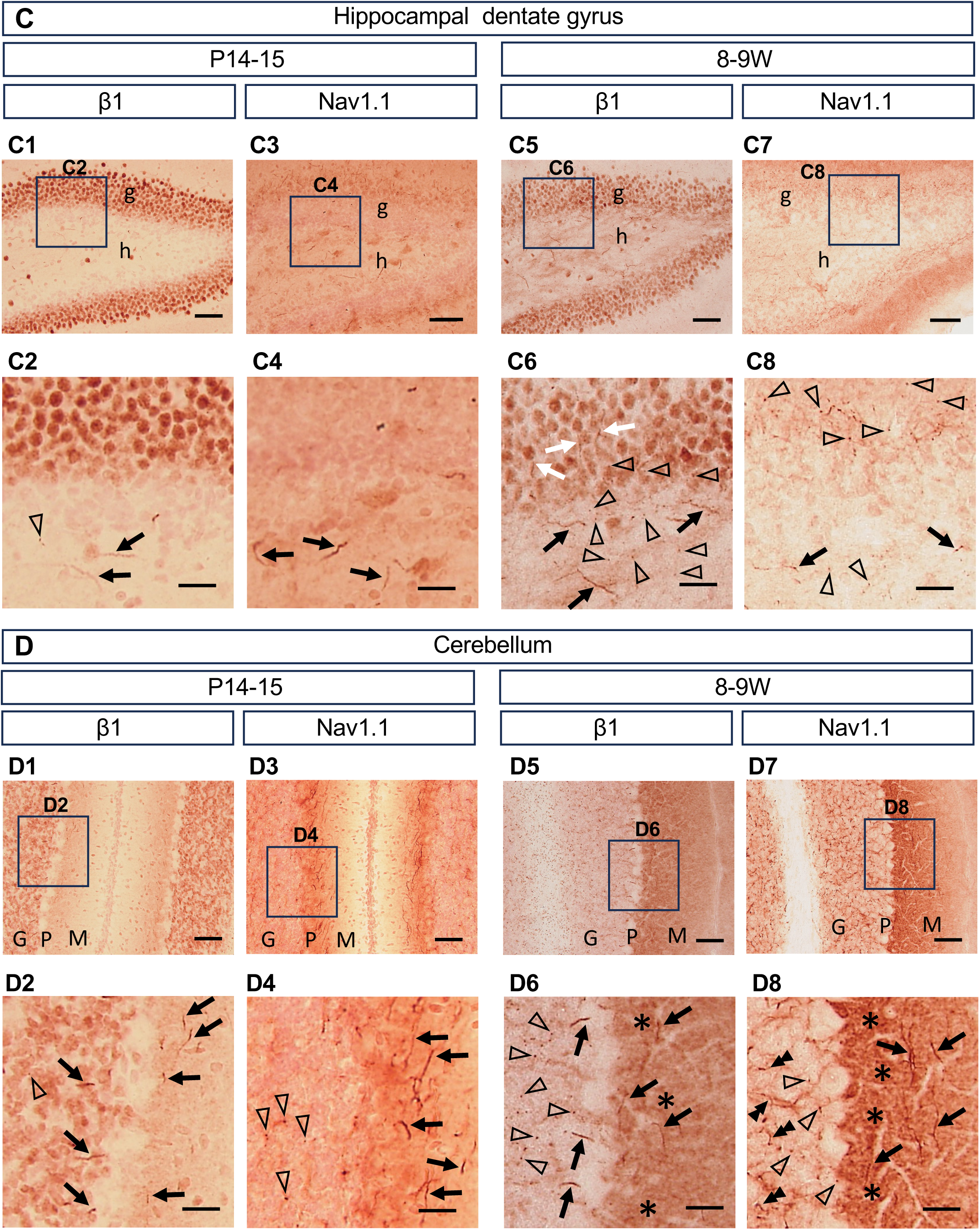
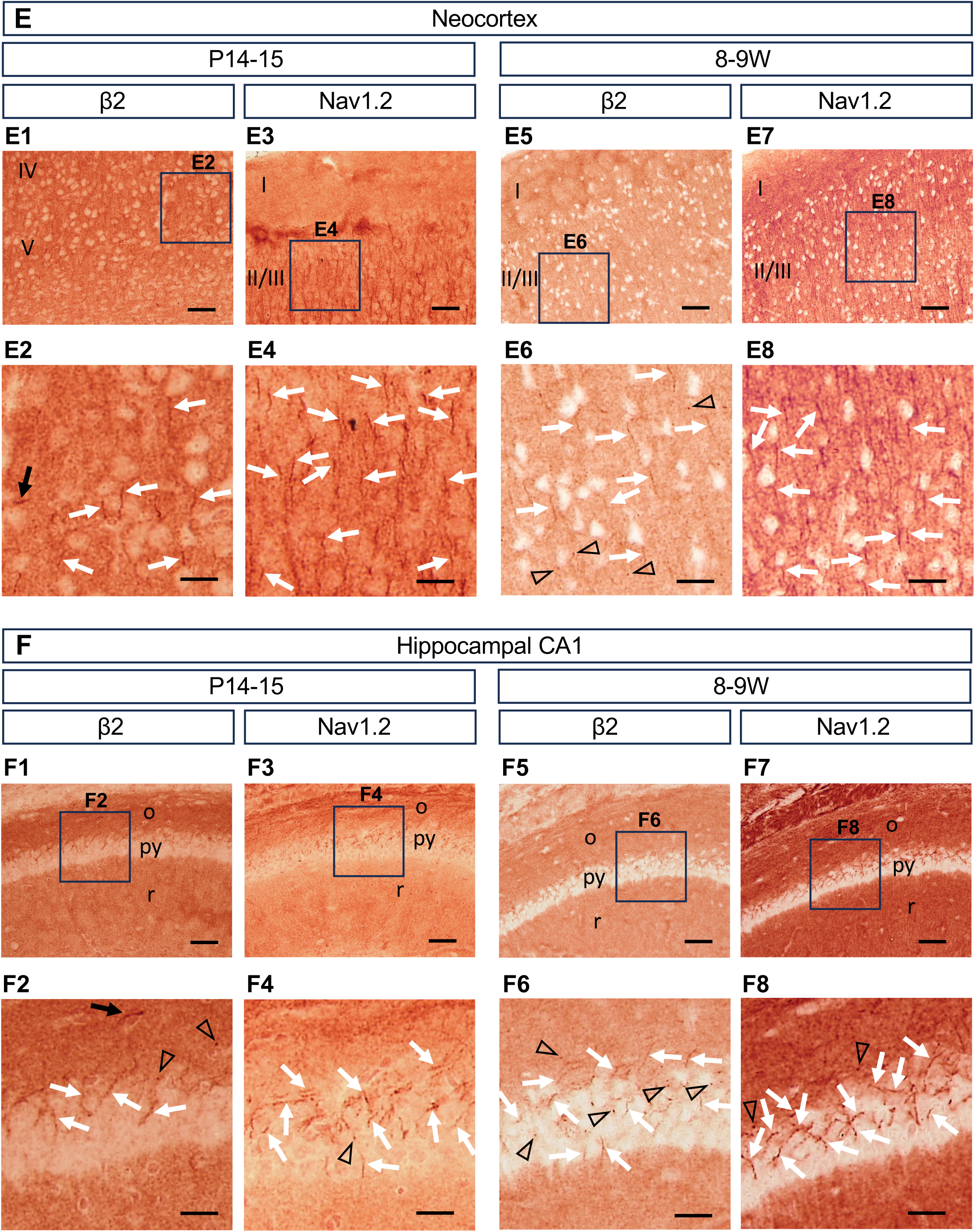
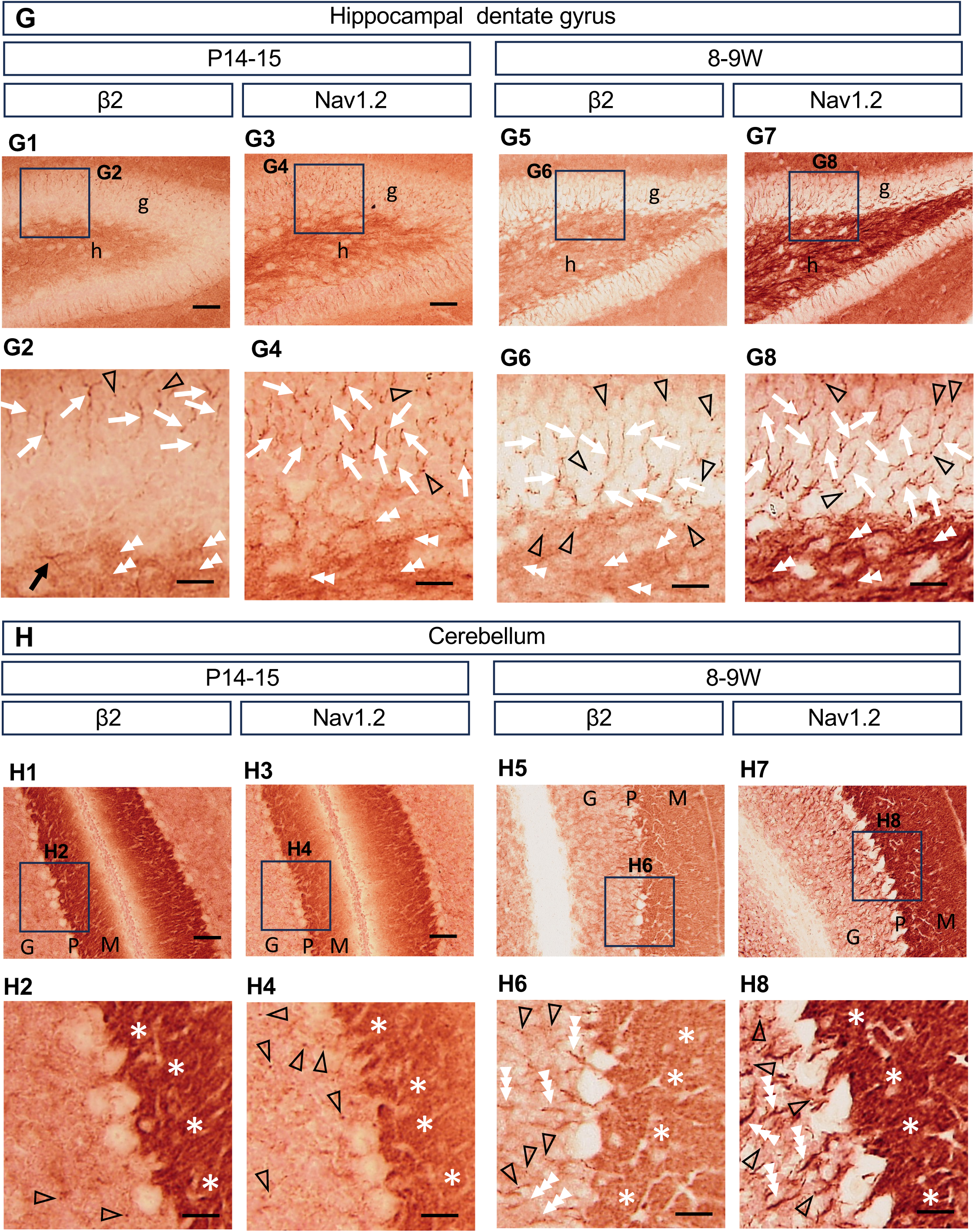
Detailed spatial distributions of the beta-1 subunit, Nav1.1, the beta-2 subunit, and Nav1.2 in the mouse brain. Chromogenic immunostaining for beta-1, Nav1.1, beta-2, or Nav1.2 was performed on parasagittal sections of C57BL/6J wild-type mouse brains at P14–15 and 8–9W, including the neocortex (A and E), hippocampus (B, C, F, and G), and cerebellum (D and H). Boxed areas in (A1, A3, A5, A7, B1, B3, B5, B7, C1, C3, C5, C7, D1, D3, D5, D7, E1, E3, E5, E7, F1, F3, F5, F7, G1, G3, G5, G7, H1, H3, H5, and H7) are shown at higher magnification in (A2, A4, A6, A8, B2, B4, B6, B8, C2, C4, C6, C8, D2, D4, D6, D8, E2, E4, E6, E8, F2, F4, F6, F8, G2, G4, G6, G8, H2, H4, H6, and H8), respectively. Arrows, arrowheads, and asterisks indicate representative signals. Black symbols indicate structures derived from inhibitory neurons, whereas white symbols indicate structures derived from excitatory neurons. Black arrows indicate the AISs of inhibitory neurons. White arrows indicate the AISs of excitatory neurons. Black double arrowheads indicate the axons of cerebellar Golgi cells. White double arrowheads indicate hippocampal mossy fibers. White triple arrowheads indicate the axons of granule cells. Open arrowheads indicate nodes of Ranvier. Black asterisks indicate distal axons of basket cells in the molecular layer of the cerebellum. White asterisks indicate parallel fibers in the molecular layer of the cerebellum. I, neocortical layer I; II/III, neocortical layers II and III; o, stratum oriens; py, stratum pyramidale; r, stratum radiatum; g, stratum granulosum; h, hilus; M, molecular layer; P, Purkinje cell layer; G, granular cell layer. Scale bars, 50 µm (A1, A3, A5, A7, B1, B3, B5, B7, C1, C3, C5, C7, D1, D3, D5, D7, E1, E3, E5, E7, F1, F3, F5, F7, G1, G3, G5, G7, H1, H3, H5, and H7) and 20 µm (A2, A4, A6, A8, B2, B4, B6, B8, C2, C4, C6, C8, D2, D4, D6, D8, E2, E4, E6, E8, F2, F4, F6, F8, G2, G4, G6, G8, H2, H4, H6, and H8).

In the neocortex at P14–15, intense beta-1 AIS signals were observed in neurons which were assumed to be inhibitory neurons based on their unparalleled directions of AISs (Fig. 2A1 and A2; black arrows), while some rather weak beta-1 signals appeared in AIS of excitatory pyramidal neurons assumed from their paralleled directions (Fig. 2A1 and A2; white arrows). In addition, a small number of punctate immunosignals of beta-1 which are assumed to be nodes of Ranvier were observed (Fig. 2A2; open arrowheads). Similar to beta-1, intense Nav1.1 signals appeared in AISs of inhibitory interneurons (Fig. 2A3 and A4; black arrows) but not much in AISs of pyramidal neurons as previously confirmed by us and other groups (see Introduction). In addition, a small number of punctate immunosignals of Nav1.1 were also observed (Fig. 2A4; open arrowheads).

In the neocortex at 8–9W, while beta-1 signal intensities at AISs of inhibitory neurons remained relatively unchanged (Fig. 2A5 and A6; black arrows), those of pyramidal neurons significantly increased (Fig. 2A5 and A6; white arrows), and the number of punctate beta-1 immunosignals also significantly increased (Fig. 2A6; open arrowheads). Similar to beta-1, Nav1.1 signals were observed in AISs of inhibitory neurons but the signal intensity was decreased (Fig. 2A7 and A8; black arrows) and instead increased in fine fibers that are assumed to be distal axons as previously reported (Ogiwara et al., 2007; Ogiwara et al., 2013). In addition, number of punctate immunosignals of Nav1.1 were significantly increased (Fig. 2A8; open arrowheads). Unlike beta-1, Nav1.1 signals were not observed in AISs of pyramidal neurons (but see Discussion).

In the hippocampal CA1 region at P14–15, intense beta-1 signals appeared in AISs of inhibitory interneurons in stratum oriens assumed from their anatomical locations (Fig. 2B1 and B2; black arrow) while weak beta-1 signals appeared in AISs of excitatory pyramidal neurons (Fig. 2B1 and B2; white arrows), and a small number of punctate immunosignals of beta-1 were observed (Fig. 2B2; open arrowheads). Similar results were also obtained in CA2–CA3 regions (Supplementary Fig. S2). Similar to beta-1, Nav1.1 was observed in AISs of inhibitory neurons at CA1–CA3 region (Fig. 2B3 and B4; black arrows), and a small number of punctate immunosignals of Nav1.1 were observed (Fig. 2B4; open arrowheads). However, unlike beta-1, Nav1.1 did not appear in pyramidal neurons (Fig. 2B3, B4 and Supplementary Fig. S2) as we previously reported (Ogiwara et al., 2007; Ogiwara et al., 2013; Yamagata et al., 2017; Yamagata et al., 2023).

In the hippocampal CA1 region at 8–9W, beta-1 signal intensity remained the same in inhibitory neurons (Fig. 2B5 and B6; black arrows), but increased significantly in pyramidal neurons (Fig. 2B5 and B6; white arrows). In addition, punctate immunosignals of beta-1 were significantly increased (Fig. 2B6; open arrowheads). On the other hand, Nav1.1 signals appeared in AISs of inhibitory neurons (Fig. 2B7 and B8; black arrow) and nodes of Ranvier (Fig. 2B8; open arrowheads). However, unlike beta-1, Nav1.1 did not appear in AISs of excitatory neurons (Fig. 2B7, B8 and Supplementary Fig. S2).

In the hippocampal dentate gyrus at P14–15, beta-1 signals appeared in AISs of inhibitory interneurons in dentate hilus assumed from their anatomical locations (Fig. 2C1 and C2; black arrows). Rarely, punctate immunosignals of beta-1 were observed (Fig. 2C2; open arrowhead). Similar to beta-1, Nav1.1 signals appeared in AISs of inhibitory neurons at dentate hilus (Fig. 2C3 and C4; black arrows). However, unlike beta-1, punctate immunosignals of Nav1.1 were not observed (Fig. 2C4).

In the hippocampal dentate gyrus at 8–9W, intense beta-1 signals were observed in AISs of inhibitory neurons in the dentate hilus (Fig. 2C5 and C6; black arrows), and beta-1–positive AISs were also observed in dentate granule cells which are excitatory neurons (Fig. 2C5 and C6; white arrows). In addition, punctate beta-1 immunosignals were significantly increased (Fig. 2C6; open arrowheads). Similar to beta-1, Nav1.1 signals were present in the AISs of inhibitory neurons in dentate hilus (Fig. 2C7 and C8; black arrows), and punctate Nav1.1 immunosignals were also significantly increased (Fig. 2C8; open arrowheads). However, in contrast to beta-1, Nav1.1 signals were not observed in dentate granule cell layer (Fig. 2C7 and C8).

In the cerebellum at P14–15, intense beta-1 signals were observed in AISs of inhibitory neurons, including Purkinje cells and basket cells, as inferred from their anatomical locations (Fig. 2D1 and D2; black arrows). In addition, a small number of punctate beta-1 immunosignals were detected (Fig. 2D2; open arrowhead). Similar to beta-1, Nav1.1 signals were also observed in the axons of basket cells (Fig. 2D3 and D4; black arrows) and in a limited number of nodes of Ranvier (Fig. 2D4; open arrowheads). However, unlike beta-1, Nav1.1 signals were not restricted to the AISs but were also detected in more distal regions of the axon (Fig. 2D3 and D4). Although previous studies have reported Nav1.1 expression in the AISs of inhibitory Purkinje cells (Van Wart and Matthews, 2006; Ogiwara et al., 2007), in the present study Nav1.1 was not detected in the AISs of Purkinje cells, possibly due to the differences in antibody affinity or experimental procedures (Fig. 2D4).

In the cerebellum at 8–9W, beta-1 signal intensities in Purkinje cells and basket cells showed no discernible changes compared with those at P14–15 (Fig. 2D5 and D6; black arrows). However, dense diffused beta-1 immunosignals appeared in the molecular layer exhibiting a decreasing gradient from the inner to the outer regions, which are assumed to be represent distal axons of basket cells (Fig. 2D5 and D6; black asterisks). In addition, punctate beta-1 immunosignals were significantly increased (Fig. 2D6; open arrowheads). Similar to beta-1, Nav1.1 signals were also detected in the AISs of presumable basket cells (Fig. 2D7 and D8; black arrows). Nav1.1 was again not detected in the AISs of Purkinje cells. Furthermore, Nav1.1 signals were detected as fibers and puncta in the granule cell layer presumably originated from inhibitory Golgi cells (Fig. 2D7 and D8; black double arrowheads). Diffuse Nav1.1 signals also increased in the molecular layer at 8–9W (Fig. 2D7 and D8; black asterisks).

These results indicate that beta-1 and Nav1.1 are predominantly co-expressed in the axons of inhibitory neurons with sole appearance of beta-1 in excitatory neurons at later developmental stage.

### 3.3. High-magnification analyses revealed preferential co-expression of beta-2 and Nav1.2 in excitatory neurons of the neocortex, hippocampus, and cerebellum

Next, we carefully compared the distributions of beta-2 and Nav1.2 at higher magnification at P14–15 and 8–9W (Fig. 2E–H), because their distributions appeared similar at low magnification (Fig. 1C and D).

In the neocortex at P14–15, intense beta-2 signals were observed in the AISs of pyramidal neurons (Fig. 2E1 and E2; white arrows) based on the parallel orientation of their AISs, as well as in some neurons presumed to be inhibitory ones based on their nonparallel orientation (Fig. 2E1 and E2; black arrow). Similar to beta-2, Nav1.2 immunosignals were densely detected in the AISs of pyramidal neurons (Fig. 2E3 and E4; white arrows). However, unlike beta-2, Nav1.2 signals were not much observed in the AISs of inhibitory neurons.

In the neocortex at 8–9W, intense beta-2 signals were more frequently observed in the AISs of pyramidal neurons (Fig. 2E5 and E6; white arrows). In contrast, the number of beta-2–positive AISs in inhibitory neurons became smaller than that at P14–15. In addition, a small number of punctate beta-2 immunosignals, presumed to represent nodes of Ranvier, were detected (Fig. 2E6; open arrowheads). Similar to beta-2, Nav1.2 immunosignals increased in the AISs of pyramidal neurons (Fig. 2E7 and E8; white arrows). Nav1.2 signals were also observed in longer fibers, which are presumed to be unmyelinated axons as described in previous studies (Yamagata et al., 2017; Ogiwara et al., 2018; Yamano et al., 2022; Yamagata et al., 2023) (Fig. 2E7 and E8).

In the hippocampal CA1 region at P14–15, beta-2 signals were observed in the AISs of excitatory pyramidal neurons (Fig. 2F1 and F2; white arrows) and in a subpopulation of inhibitory neurons (Fig. 2F1 and F2; black arrow). In addition, a small number of punctate beta-2 immunosignals were observed (Fig. 2F2; open arrowheads). Similar to beta-2, intense Nav1.2 signals were observed in the AISs of pyramidal neurons (Fig. 2F3, F4; white arrows, and Supplementary Fig. S2), as well as in a few nodes of Ranvier (Fig. 2F4; open arrowhead). However, unlike beta-2, Nav1.2 was not observed in inhibitory neurons.

In the hippocampal CA1 region at 8–9W, beta-2 signals increased in the AISs and in some nodes of Ranvier within the pyramidal cell layer (Fig. 2 F6; white arrows and open arrowheads). Similar to beta-2, Nav1.2 signals also increased in the AISs of pyramidal neurons (Fig. 2F7, F8; white arrows, and Supplementary Fig. S2), as well as in a few nodes of Ranvier (Fig. 2F8; open arrowheads). However, unlike beta-2, diffuse Nav1.2 immunosignals in layers of oriens and radiatum became much denser at 8–9W than those at P14–15.

In the hippocampal dentate gyrus at P14–15, beta-2 signals were observed along the excitatory mossy fibers (Fig. 2G1 and G2; white double arrowheads). In addition, beta-2 signals were detected in some AISs of inhibitory neurons in the dentate hilus (Fig. 2G1 and G2; black arrow), as well as in a few nodes of Ranvier (Fig. 2G2; open arrowheads). Similar to beta-2, intense Nav1.2 immunosignals were observed in the AISs of dentate granule cells (Fig. 2G3 and G4; white arrows), in mossy fibers (Fig. 2G3 and G4; white double arrowheads), and in nodes of Ranvier (Fig. 2G4; open arrowheads). However, unlike beta-2, Nav1.2 signals were not detected in the AISs of inhibitory neurons in the dentate hilus (Fig. 2G3 and G4).

In the hippocampal dentate gyrus at 8–9W, the localization of beta-2 signals in the AISs of excitatory granule cells (Fig. 2G5 and G6; white arrows) and in mossy fibers (Fig. 2G5 and G6; white double arrowheads) was unchanged compared with that at P14–15, but their signal intensities were increased. The number of punctate beta-2 signals also increased significantly (Fig. 2G6; open arrowheads). In contrast, the number of beta-2–positive AISs in inhibitory neurons decreased at 8–9W compared with P14–15, as also observed in the neocortex. Similar to beta-2, Nav1.2 immunosignal intensity was increased at the AISs of granule cells (Fig. 2G7 and G8; white arrows) and mossy fibers (Fig. 2G7 and G8; white double arrowheads) as well as at nodes of Ranvier (Fig. 2G8; open arrowheads) compared with P14–15, which is consistent with previous studies (Liao et al., 2010; Ogiwara et al., 2018).

In the cerebellum at P14–15, beta-2 signals were broadly and densely distributed throughout the molecular layer, presumably the parallel fibers of granule cells which are the only excitatory neurons of the cerebellum based on their anatomical characteristics (Fig. 2H1 and H2; white asterisks). In addition, a small number of punctate beta-2 immunosignals were observed in the granular layer (Fig. 2H2; open arrowheads). Similar to beta-2, defused Nav1.2 signals were densely detected in the molecular layer (Fig. 2H3 and H4; white asterisks). Some punctate Nav1.2 immunosignals were also observed in the granular layer (Fig. 2H4; open arrowheads).

In the cerebellum at 8–9W, beta-2 signals were observed in the presumed proximal (Fig. 2H5 and H6; white triple arrowheads) and distal axons of granule cells (Fig. 2H5 and H6; white asterisks), as well as at nodes of Ranvier (Fig. 2H6; open arrowheads). No beta-2 immunosignals were detected in cerebellar Purkinje cells. Similar to beta-2, Nav1.2 immunosignals were localized to the distal axons of granule cells (i.e., parallel fibers) and were uniformly distributed throughout the molecular layer (Fig. 2H7 and H8; white asterisks). In addition, intense Nav1.2 immunosignal appeared in the presumed proximal axons of granule cells (Fig. 2H7 and H8; white triple arrowheads) and at nodes of Ranvier (Fig. 2H8; open arrowheads). The homogeneous Nav1.2 staining showed a high degree of correspondence with the beta-2 staining pattern in the molecular layer. In contrast, the homogeneous staining patterns of beta-2 and Nav1.2 differed from the gradient-like distributions observed for Nav1.1 and beta-1. These findings indicate that cerebellar granule cells express Nav1.2 and beta-2, but not Nav1.1 and beta-1.

These results indicate that beta-2 and Nav1.2 are predominantly localized to the AISs of excitatory neurons and that their expression increases in unmyelinated axons, such as mossy fibers, during development.

### 3.4. Nav1.6 is broadly expressed at the AISs of neocortical, hippocampal, and cerebellar neurons and remains proximally localized at adult stages

In the low-magnification analysis, the distribution of Nav1.6 differed from that of Nav1.1 and Nav1.2. However, co-localization of Nav1.6 with Nav1.1 or Nav1.2 has been reported in neocortical interneurons and hippocampal pyramidal neurons, respectively (Lorincz and Nusser, 2008; Lorincz and Nusser, 2010). Therefore, we further examined the distribution of Nav1.6 at higher magnification.

In the neocortex at P14–15, Nav1.6 signals were observed in the AISs of a subpopulation of pyramidal neurons (Fig. 3A1 and A2; white arrows) and inhibitory neurons (Fig. 3A1 and A2; black arrows). Nav1.6 signals were more intense in the AISs of inhibitory neurons than in those of pyramidal neurons. The distribution pattern of Nav1.6 was more similar to that of beta-1 than to that of beta-2.

**Figure 3.**
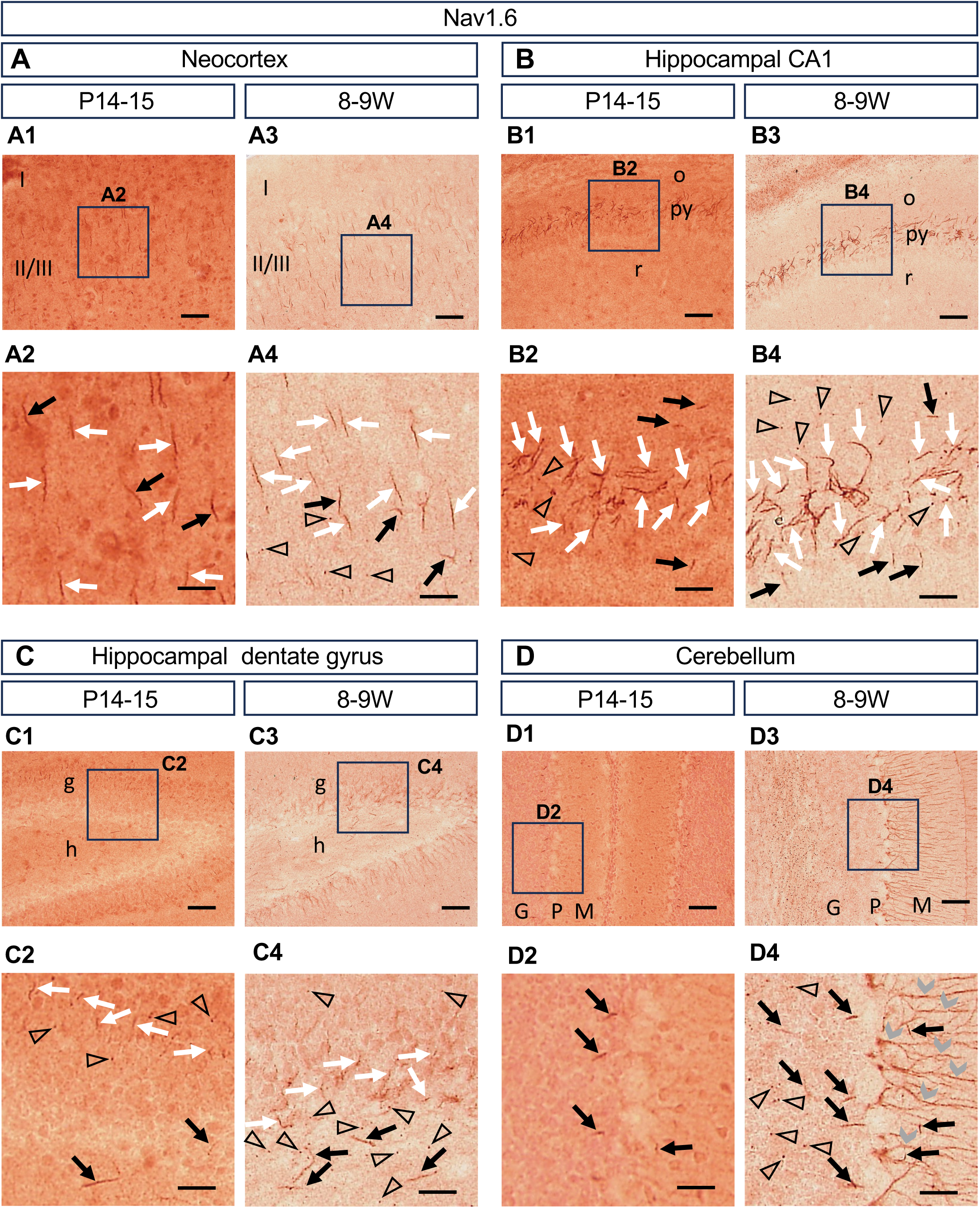
Nav1.6 distribution largely overlaps with that of Nav1.1 or Nav1.2 in the mouse brain. Chromogenic immunostaining for Nav1.6 was performed on parasagittal sections of C57BL/6J wild-type mouse brains at P14–15 and 8–9W, including the neocortex (A), hippocampus (B and C), and cerebellum (D). Magnified images outlined in the upper panels (A1, A3, B1, B3, C1, C3, D1, and D3) are shown in the lower panels (A2, A4, B2, B4, C2, C4, D2, and D4), respectively. Arrows, arrowheads, and wedges indicate representative signals. Black arrows indicate the AISs of inhibitory neurons. White arrows indicate the AISs of excitatory neurons. Gray wedges indicate Bergmann glial fibers. Open arrowheads indicate nodes of Ranvier. I, neocortical layer I; II/III, neocortical layers II and III; o, stratum oriens; py, stratum pyramidale; r, stratum radiatum; g, stratum granulosum; h, hilus; M, molecular layer; P, Purkinje cell layer; G, granular cell layer. Scale bars, 50 µm (A1, A3, B1, B3, C1, C3, D1, and D3) and 20 µm (A2, A4, B2, B4, C2, C4, D2, and D4).

In the neocortex at 8–9W, intense Nav1.6 signals were more frequently observed in the AISs of pyramidal neurons (Fig. 3A3 and A4; white arrows) and inhibitory neurons (Fig. 3A3 and A4; black arrows) than at P14–15. In addition, a small number of punctate Nav1.6 immunosignals, presumed to represent nodes of Ranvier, were present (Fig. 3A4; open arrowheads). Similar to P14–15, the distribution patterns of Nav1.6 and beta-1 were comparable.

In the hippocampal CA1 region at P14–15, Nav1.6 signals were observed in the AISs of the majority of pyramidal neurons (Fig. 3B1 and B2; white arrows), a subset of inhibitory neurons (Fig. 3B1 and B2; black arrows), and nodes of Ranvier (Fig. 3B2; open arrowheads), consistent with previous studies (Liao et al., 2010; Ogiwara et al., 2018). A similar distribution pattern of Nav1.6 was observed in the CA2–CA3 regions (Supplementary Fig. S2). The distribution pattern of Nav1.6 closely resembled that of beta-2/Nav1.2 in excitatory neurons and that of beta-1/Nav1.1 in inhibitory neurons.

In the hippocampal CA1 region at 8–9W, Nav1.6 signals were observed in the AISs of pyramidal neurons (Fig. 3B3 and B4; white arrows) and inhibitory neurons (Fig. 3B3 and B4; black arrows), with stronger signal intensity than at P14–15. In addition, punctate Nav1.6 immunosignals were significantly increased (Fig. 3B4; open arrowheads). Similar to P14–15, the distribution patterns of Nav1.6 and beta-1 were comparable.

In the hippocampal dentate gyrus at P14–15, Nav1.6 signals were observed in the AISs of granule cells (Fig. 3C1 and C2; white arrows) and nodes of Ranvier (Fig. 3C2; open arrowheads). In the dentate hilus, Nav1.6 signals appeared in some inhibitory neurons (Fig. 3C1 and C2; black arrows). The distribution of Nav1.6 was similar to that of beta-1/Nav1.1 in the dentate hilus and to that of beta-2/Nav1.2 in the granule cell layer. However, unlike beta-2 and Nav1.2, Nav1.6 signals were not detected in mossy fibers.

In the hippocampal dentate gyrus at 8–9W, Nav1.6 signals increased in the AISs of granule cells (Fig. 3C3 and C4; white arrows) and inhibitory neurons (Fig. 3C3 and C4; black arrows), as well as in nodes of Ranvier (Fig. 3C4; open arrowheads), compared with P14–15. Similar to P14–15, Nav1.6 and beta-1/Nav1.1 showed comparable distribution patterns in the dentate hilus, whereas Nav1.6 and beta-2/Nav1.2 exhibited similar distributions in the granule cell layer.

In the cerebellum at P14–15, Nav1.6 signals were observed in the AISs of Purkinje cells and basket cells (Fig. 3D1 and D2; black arrows), similar to beta-1 signals. In the molecular layer, Nav1.6 immunosignals did not exhibit a diffuse distribution pattern, in contrast to those of beta-1/Nav1.1 or beta-2/Nav1.2.

In the cerebellum at 8–9W, Nav1.6 immunosignals were observed in the AISs of Purkinje cells, basket cells, and a subset of AISs of neurons presumed to be Golgi cells, based on their anatomical locations (Fig. 3D3 and D4; black arrows), as well as at nodes of Ranvier (Fig. 3D4; open arrowheads). In addition, Nav1.6 immunosignals were observed in radial fibers within the molecular layer that are morphologically consistent with Bergmann glial processes (Fig. 3D3 and D4; gray wedges), as previously reported (Caldwell et al., 2000; Krzemien et al., 2000). Similar to P14–15, Nav1.6 and beta-1 showed comparable distribution patterns in Purkinje cells and basket cells, but the comparison in Golgi cells remains unclear.

These results indicate that Nav1.6 is expressed at the AISs of inhibitory and excitatory neurons at the juvenile stage (P14–15). At the later adult stage (8–9W), Nav1.6 expression in these neurons is further increased.

Taken together with the results for beta-1, beta-2, Nav1.1, and Nav1.2 described above, these findings suggest that Nav1.6 may be co-expressed with beta-1 and Nav1.1 in subsets of inhibitory neurons and with beta-1, beta-2, and Nav1.2 in subsets of excitatory neurons.

## 4. Discussion

In this study, low-magnification analyses revealed that the distributions of beta-1 and beta-2 subunits in the mouse brain closely resemble those of Nav1.1 and Nav1.2 subunits, respectively. Beta-1 and Nav1.1 were predominantly distributed in caudal regions, including the thalamus, hypothalamus, midbrain, cerebellum, and medulla oblongata, whereas beta-2 and Nav1.2 were mainly localized to rostral regions such as the neocortex, hippocampus, and striatum. In contrast, Nav1.6 did not exhibit a clear rostrocaudal bias. High-magnification analyses further demonstrated that the three alpha-subunits (Nav1.1, Nav1.2, and Nav1.6) and the two beta-subunits (beta-1 and beta-2) were primarily localized to the AIS and/or nodes of Ranvier. Together, these spatial distributions of VGSC alpha– and beta-subunits suggest distinct combinatorial patterns in the mouse brain.

Our previous study demonstrated that the expression of Nav1.1 and Nav1.2 is mutually exclusive (Yamagata et al., 2017). Subsequent analyses revealed that Nav1.1 is selectively expressed in inhibitory neurons in the hippocampus, whereas in the cortex it is expressed not only in inhibitory neurons but also in a subset of excitatory neurons, including some corticocortical projection neurons and pyramidal tract projection neurons (Yamagata et al., 2023). Although Nav1.1 expression in neocortical excitatory neurons was not detected in the present study, likely because of its low expression level, integrating these findings supports the possibility of physiologically relevant combinations of Nav1.1, Nav1.2, Nav1.6, beta-1, and beta-2 in distinct inhibitory and excitatory neuronal populations in the mouse brain. These combinatorial expression patterns are schematically summarized in Fig. 4.

**Figure 4.**
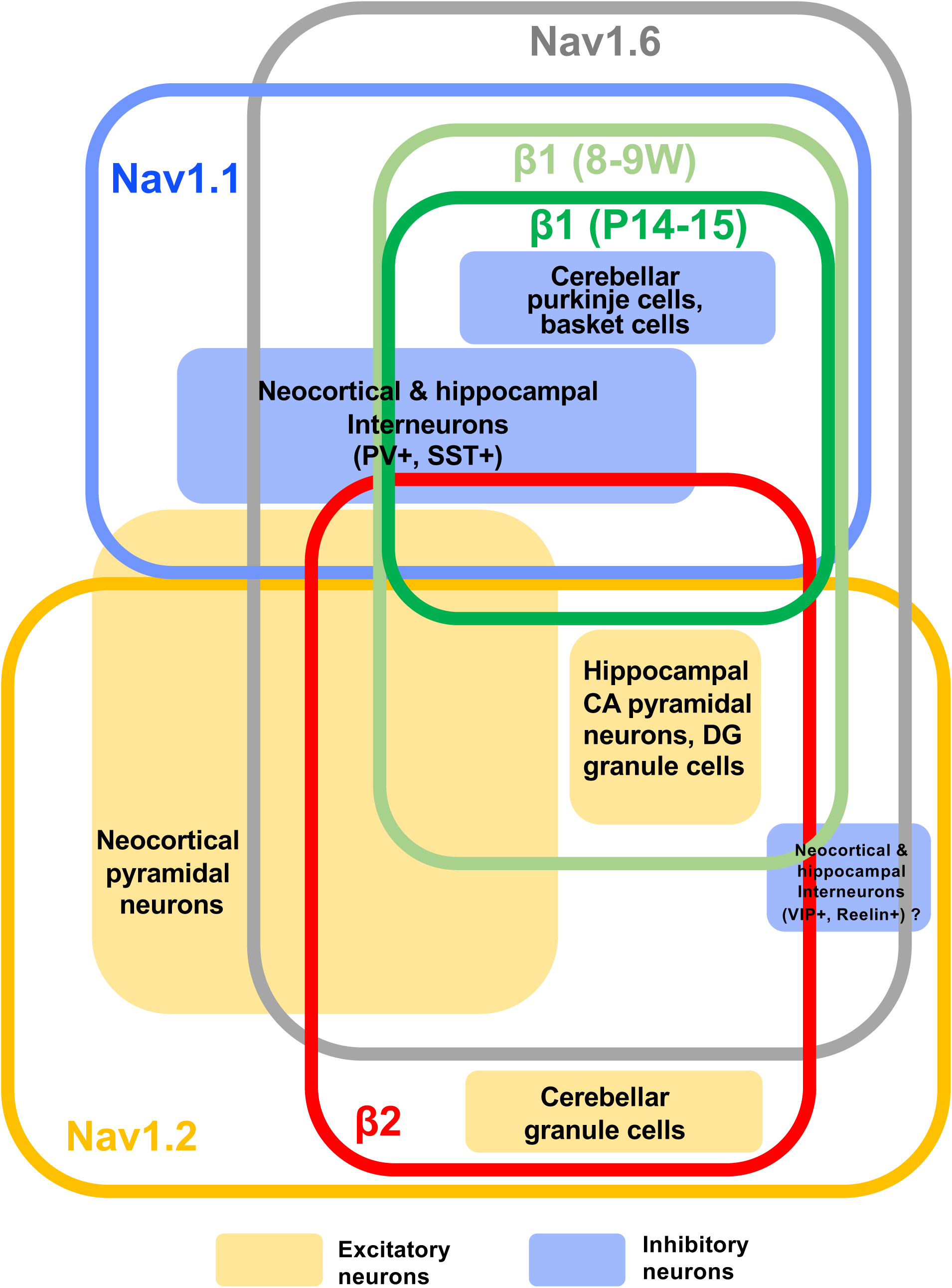
Expression profiles of VGSC subunits in excitatory and inhibitory neurons. Schematic summary of putative combinatorial expression patterns of VGSC alpha and beta subunits in inhibitory and excitatory neurons based on the present histological analyses. The expression of Nav1.1 and Nav1.2 is largely mutually exclusive. The expression of Nav1.6 is broadly distributed in both excitatory and inhibitory neurons. The beta-1 subunit is expressed in both Nav1.1–positive inhibitory neurons and Nav1.2–positive excitatory neurons, whereas the beta-2 subunit is primarily expressed in Nav1.2–positive excitatory neurons. Excitatory and inhibitory neurons are indicated by light orange and light blue boxes, respectively.

Although beta-1 has been reported to associate noncovalently with alpha subunits in vivo (Hartshorne et al., 1982), the specific brain regions in which it interacts with Nav1.1, Nav1.2, and Nav1.6 remain unclear. In the present study, we found that beta-1 is predominantly localized in the AISs of inhibitory neurons in the neocortex, hippocampus, and cerebellum, particularly during juvenile stages. Beta-1 is also localized in the AIS of excitatory neurons, and its immunoreactivity increased during adult stages. These results are consistent with those reported by Wimmer et al. (2015), which demonstrated beta-1 subunit expression at the AIS of both inhibitory and excitatory neurons in the cerebral cortex, hippocampus, and cerebellum. At juvenile stages, the expression pattern of beta-1 closely resembled that of Nav1.1 in the neocortex, hippocampus, and cerebellum. Based on these findings and previous studies of *Scn1a*-deficient mice (Ogiwara et al., 2013; Tatsukawa et al., 2018) and *Scn1b*-deficient mice (Chen et al., 2004), we hypothesize that loss of function of either beta-1 or Nav1.1 during early development may contribute to overlapping neurological phenotypes through dysfunction of inhibitory neurons. This hypothesis is supported by reports demonstrating amelioration of symptoms in Nav1.1-deficient mice following expression of a beta-1 transgene (Niibori et al., 2020), as well as by the identification of *SCN1B* mutations in a subset of patients with Dravet syndrome (Patino et al., 2009; Ogiwara et al., 2012) and in patients with GEFS+ (Wallace et al., 1998; Scheffer et al., 2007) in the absence of *SCN1A* mutations.

With respect to beta-2, it was primarily detected at the AIS of excitatory neurons, and strong immunoreactivity was also observed in unmyelinated axons, similar to that of Nav1.2 in the neocortex, hippocampus, and cerebellum. Although these results differ from the beta-2 localization to cell bodies and dendrites in the rat neocortex, hippocampus, and cerebellum reported by Yu et al. (2003), this discrepancy is likely attributable not to species differences but rather to differences in antibody specificity and immunostaining methodologies. In a previous study of *Scn2b*^−/−^ mice, the loss of beta-2 in these brain regions did not result in spontaneous seizures or sudden unexpected death in epilepsy, nor did it affect beta-1 protein levels or the distribution of Nav1.6 at nodes of Ranvier in the sciatic and optic nerve (Chen et al., 2002). However, *Scn2b*^−/−^ mice exhibited reduced VGSC expression at the plasma membrane and increased sensitivity to pilocarpine. In contrast, mutations in *SCN2A* have been identified as causes of epilepsy, ASD, and ID (reviewed in Yamakawa, 2016; Meisler et al., 2021). The broad phenotypic spectrum associated with *SCN2A* mutations suggests the involvement of modifier genes. Given that the beta-2 subunit is covalently linked to the VGSC alpha subunit (Hartshorne et al., 1982), our findings suggest that beta-2 may act as a modifier of the phenotypic consequences of *SCN2A* mutations, potentially contributing to variability in epilepsy, ASD, and ID.

Furthermore, our results show that Nav1.6 is expressed in both inhibitory and excitatory neurons, exhibiting a distribution pattern that overlaps with those of beta-1 and Nav1.1 or beta-2 and Nav1.2. Numerous de novo mutations in *SCN8A* have been identified as causes of severe epileptic encephalopathy (reviewed in Yamakawa et al., 2024). The observation that de novo mutations in *SCN2A* also lead to severe epileptic encephalopathy, together with the abundant expression of Nav1.6 in excitatory neurons, suggests that dysfunction of neocortical excitatory neurons may underlie severe epileptic encephalopathy. Conversely, although reduced *Scn8a* expression induces seizures in mice, it has also been shown to ameliorate phenotypes associated with Nav1.1 deficiency (Makinson et al., 2016; Lenk et al., 2020). These findings suggest that impairment of inhibitory neurons expressing Nav1.1 may be counteracted by reduced function in excitatory neurons expressing Nav1.6. Collectively, these results raise the possibility that partial functional reduction of Nav1.6 or Nav1.2 in excitatory neurons could represent a therapeutic strategy for Dravet syndrome. Because this study is based on histological analyses, direct functional interactions among VGSC subunits and their causal contributions to disease phenotypes remain to be determined.

Recently, advances in image-based in situ mRNA quantification techniques, such as in situ sequencing and single-cell resolution in situ hybridization on tissues (SCRINSHOT), have enabled high-throughput analysis of gene expression in tissues (Sountoulidis et al., 2020). While these approaches offer high sensitivity for mRNA detection, they provide limited information regarding subcellular localization. In contrast, conventional immunohistochemical staining has lower detection sensitivity, which depends on the properties of both target proteins and antibodies, but allows direct visualization of protein localization and colocalization within tissues. Thus, classical immunohistochemical approaches remain valuable tools for uncovering novel biological insights.

In summary, this study provides a comprehensive comparison of VGSC subunit expression in neurons using immunohistochemical approaches. These findings may help define brain regions and neural circuit alterations that contribute to epilepsies and neurodevelopmental disorders caused by mutations in VGSC subunit genes.

## CRediT authorship contribution statement

**Tetsushi Yamagata:** Conceptualization, Data curation, Validation, Writing – original draft, Writing – review & editing, Funding acquisition. **Toshimitsu Suzuki:** Data curation, Validation, Writing – original draft, Writing – review & editing. **Kazuhiro Yamakawa:** Conceptualization, Funding acquisition, Project administration, Supervision, Writing – original draft, Writing – review & editing.

## Acknowledgements

We are grateful to all members of the Department of Neurodevelopmental Disorder Genetics at NCU and the Laboratory for Neurogenetics at RIKEN CBS for helpful discussions. We also thank Drs. Fumitaka Oyama and Nobuyuki Nukina (RIKEN CBS) for providing the anti-beta-1 and anti-beta-2 antibodies.

## Supplementary figure legends

**Supplementary Figure S1. A web-based database for the distributions of VGSC subunits in the mouse brain.** Immunostaining images of VGSC subunits are available on our website (https://ndg-ibs.github.io/VGSC_IHC_DB/).

**Supplementary Figure S2. The distribution of Nav1.6 largely overlaps with that of Nav1.1 or Nav1.2 in hippocampal CA2 and CA3.** Chromogenic immunostaining for Nav1.6 was performed on parasagittal sections from C57BL/6J mouse brains at P14–15 or 8–9W. Magnified images outlined in each region (left panels) are shown in the right panels. o, stratum oriens; py, stratum pyramidale; r, stratum radiatum; l, stratum lucidum; g, stratum granulosum; h, hilus; M, molecular layer; P, Purkinje cell layer; G, granular cell layer. Scale bars: 50 µm (low magnification) and 20 µm (high magnification).

## Highlights

∎ Combinations of VGSC subunits are largely unique among neuronal subpopulations.
∎ Beta-1 is predominantly expressed in inhibitory neurons in early development.
∎ Beta-2 is preferentially expressed in Nav1.2-positive excitatory neurons.

