## Supplementary figures for "Parallel analysis of voltage-gated sodium channel subunits reveals preferential colocalizations of beta-1/Nav1.1 and beta-2/Nav1.2"

### Distribution of VGSC Subunits in Mouse Brain (strain: C57BL/6J)

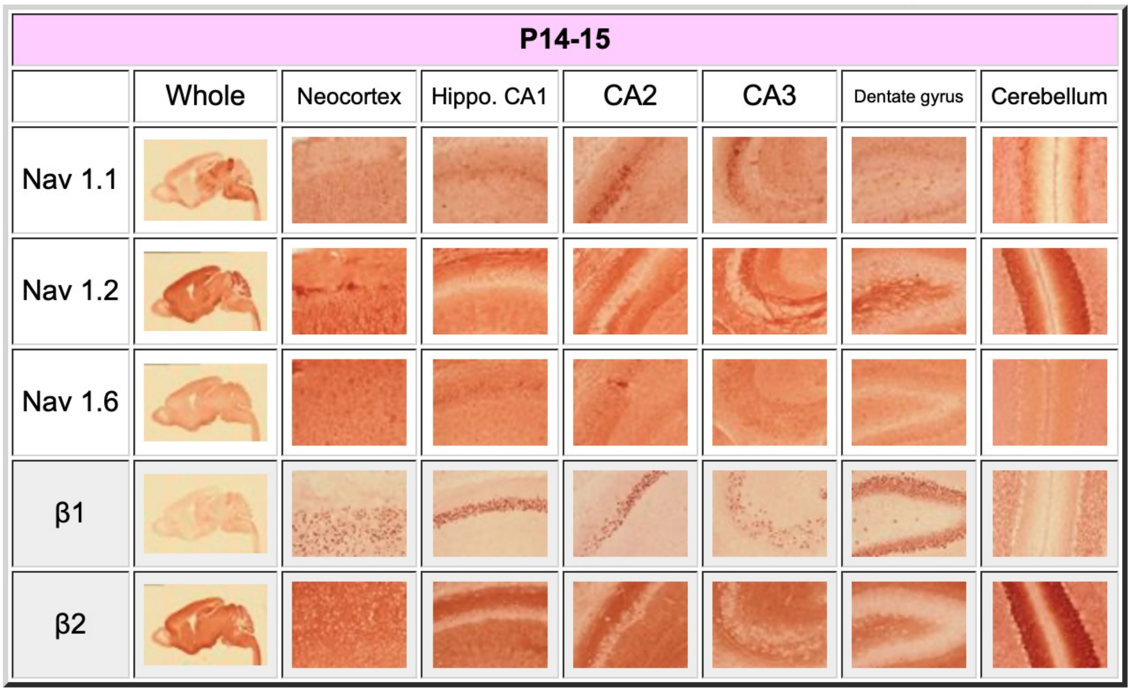


### Distribution of VGSC Subunits in Mouse Brain (strain: C57BL/6J)

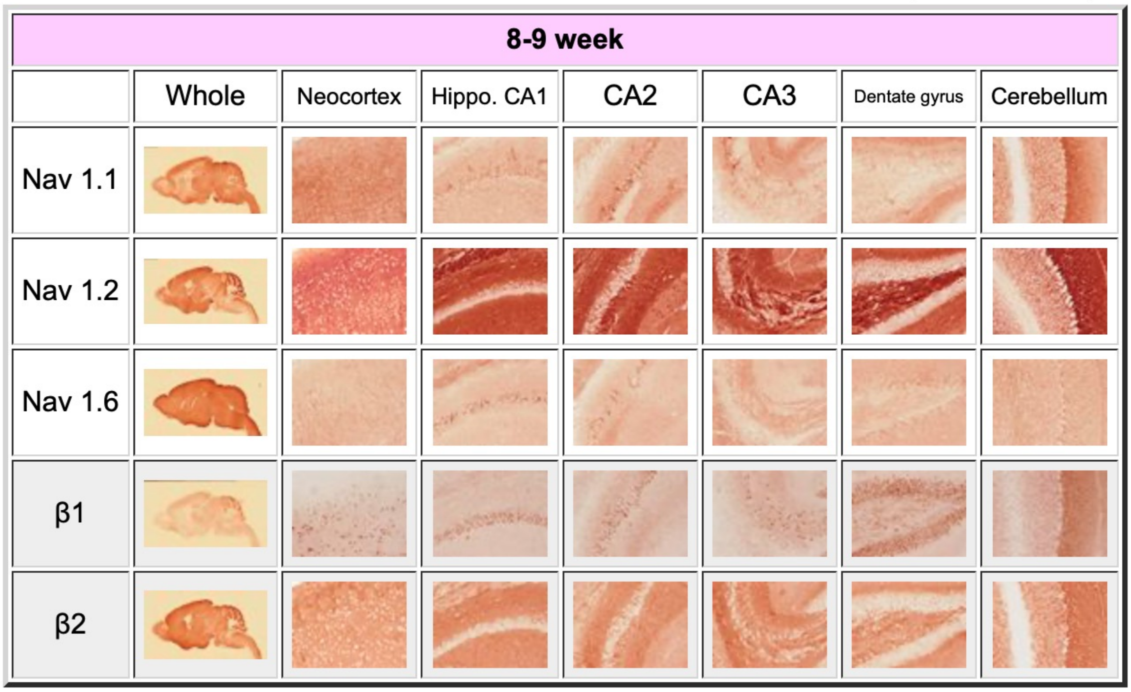


Yamagata et al., Supplementary figure S1

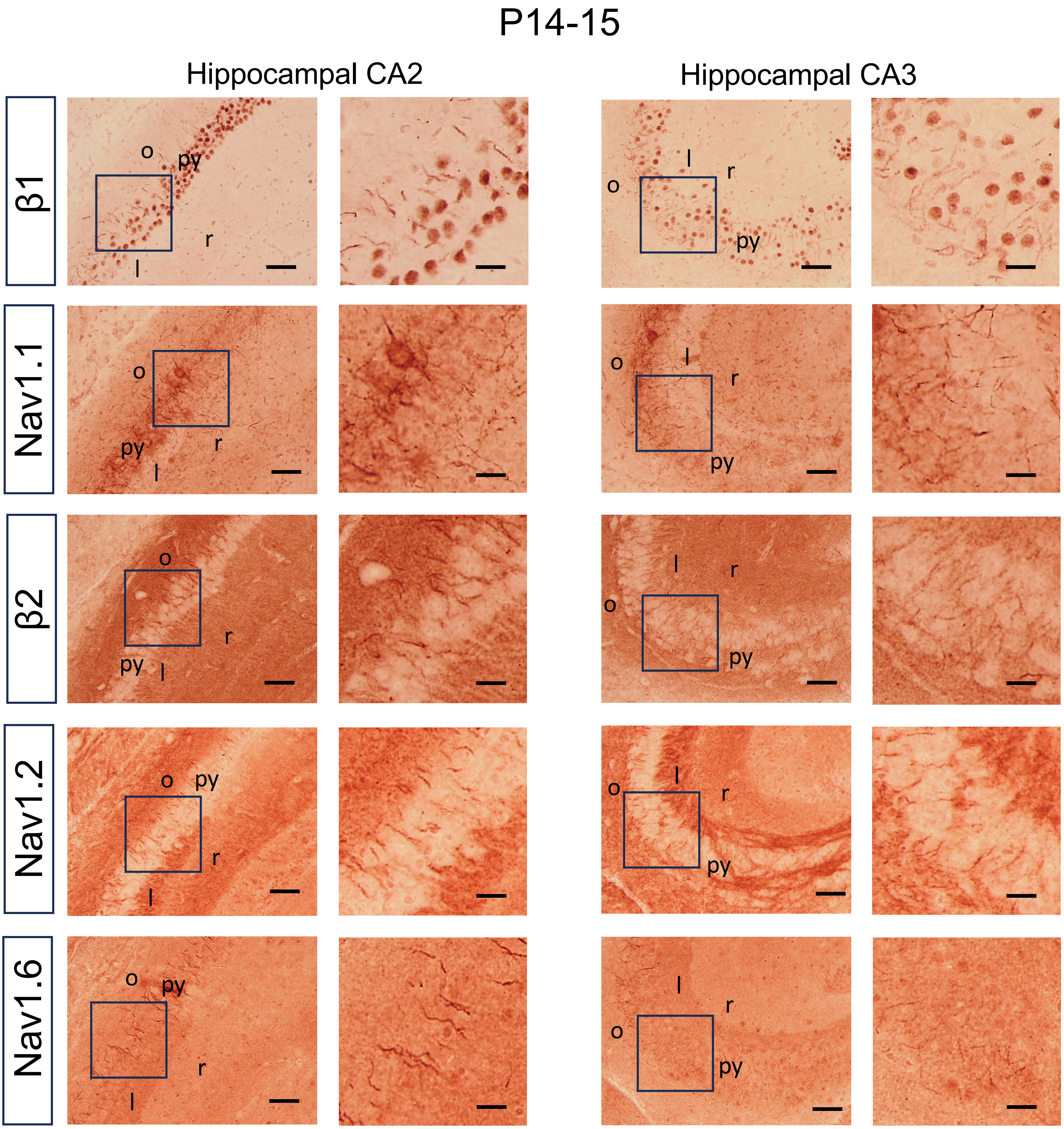


Yamagata et al., Supplementary figure S2

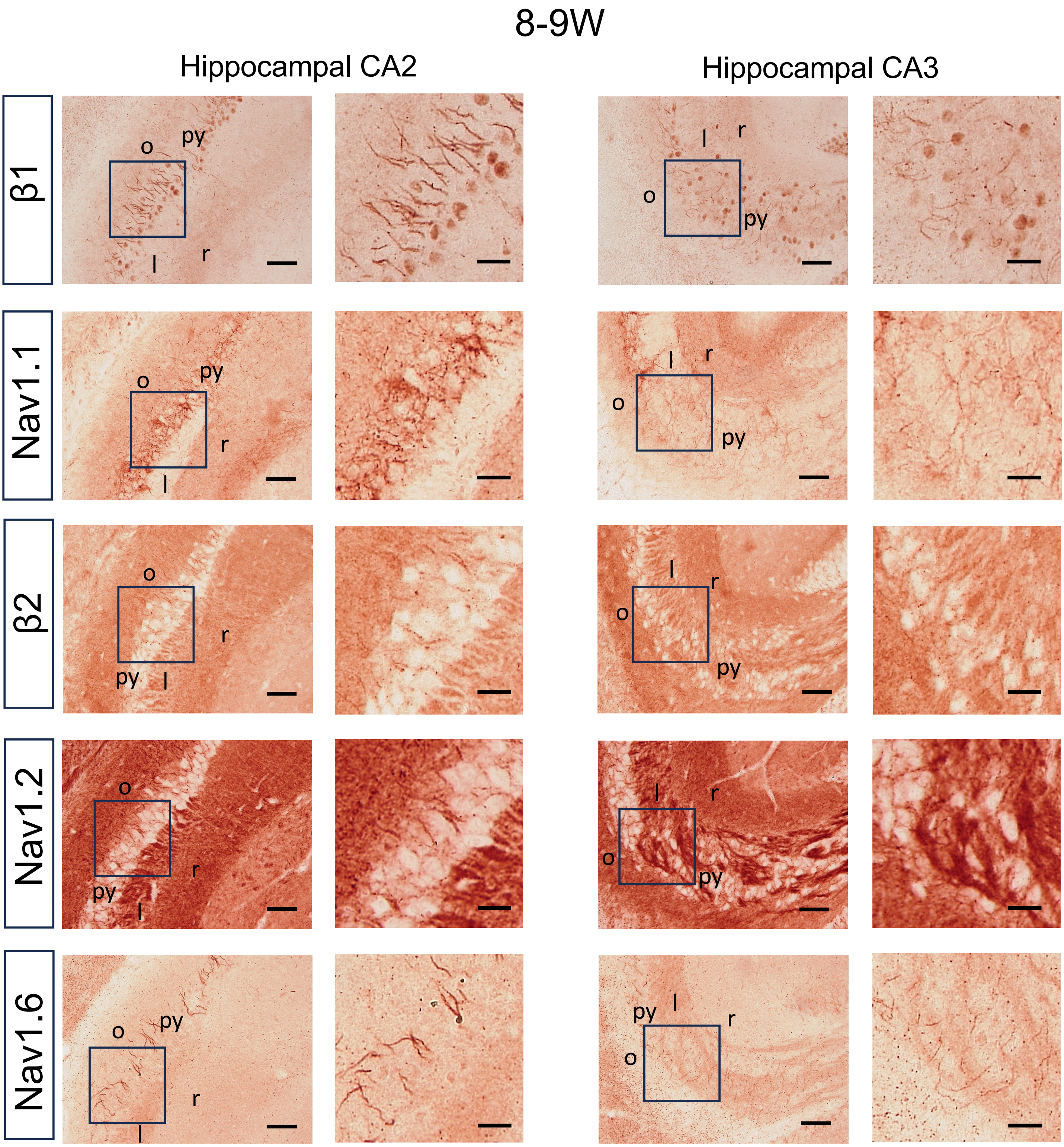


Yamagata et al., Supplementary figure S2 Continued
